# Comparative analysis of *Streptococcus* genomes

**DOI:** 10.1101/447938

**Authors:** Pavel V Shelyakin, Olga O Bochkareva, Anna A Karan, Mikhail S Gelfand

**Affiliations:** Vavilov Institute of General Genetics Russian Academy of Sciences, Moscow, Russia; Kharkevich Institute for Information Transmission Problems, Moscow, Russia; Center for Data-Intensive Biomedicine and Biotechnology, Skolkovo Institute of Science and Technology, Moscow, Russia; Faculty of Bioengineering and Bioinformatics, Lomonosov Moscow State University; Faculty of Computer Science, Higher School of Economics, Moscow, Russia

**Author notes:** equal contribution (Pavel V Shelyakin, Olga O Bochkareva).

**Keywords:** *Streptococcus*, genome rearrangements, pan-genome, antigen variation, gene inflow, selection in upstream regions

## Abstract

**Background:** Genome sequencing of multiple strains demonstrated high variability in gene content even in closely related strains of the same species and created a newly emerged object for genomic analysis, the pan-genome, that is, the complete set of genes observed in a given species or a higher level taxon. Here we analysed the pan-genome structure and the genome evolution of 25 strains of *Streptococcus suis*, 50 strains of *Streptococcus pyogenes* and 28 strains of *Streptococcus pneumoniae*.

**Results:** Fractions of the pan-genome, unique, periphery, and universal genes differ in size, functional composition, the level of nucleotide substitutions, and predisposition to horizontal gene transfer and genomic rearrangements. The density of substitutions in intergenic regions appears to be correlated with selection acting on adjacent genes, implying that more conserved genes tend to have more conserved regulatory regions. The total pan-genome of the genus is open, but only due to strain-specific genes, whereas other pan-genome fractions reach saturation. The strain-specific fraction is enriched with mobile elements and hypothetical proteins, but also contains a number of candidate virulence-related genes, so it may have a strong impact on adaptability and pathogenicity.

About 7% of single-copy periphery genes have been found in different syntenic regions. More than a half of these genes are rare in all *Streptococcus* species; others are rare in at least one species. We have identified the set of genes with phylogenies inconsistent with species and non-conserved location in the chromosome; these genes are candidates for horizontal transfer between species.

An inversion of length 15 kB found in four independent branches of *S. pneumoniae* has breakpoints formed by genes encoding a surface antigen protein (PhtD). The observed parallelism may indicate the action of an antigen variation mechanism.

**Conclusions:** Members of the genus *Streptococcus* have a highly dynamic, open pan-genome, that potentially confers them with the ability to adapt to changing environmental conditions, i.e. antibiotic resistance or transmission between different hosts. Hence, understanding of genome evolution is important for the identification of potential pathogens and design of drugs and vaccines.

## Background

The genus *Streptococcus* are Gram-positive bacteria that exert strong influence on the health of humans and animals. In particular, *Streptococcus pneumoniae*, normally a commensal from the nasopharynx microflora, at the same time is responsible for most pneumonia cases and is second only to *Mycobacterium tuberculosis* as a cause of mortality from bacterial infection worldwide (Krzyściak et al., 2013). *Streptococcus pyogenes* is among the top ten of bacterial causes of human mortality worldwide (Brown, Gilliland, and Holden, 2001; Richards et al., 2014), and due to the molecular mimicry with heart and brain cells causes severe autoimmune sequelae like rheumatic fever (Ferretti, Stevens, and Fischetti, 2016) and, possibly, autoimmune neuropsychiatric disorders (Mullen, 2015). *Streptococcus suis* rarely causes disease in human, but is one of the most important swine pathogens (Gottschalk and Segura, 2000).

Sequencing of multiple strains of one species has demonstrated that the genome of any single strain does not reflect the genetic variability of the species, as two strains may differ by 20–35% of the gene content (Medini et al., 2005). The concept of pan-genome was introduced to represent the total set of genes observed in genomes of strains assigned to a given species (Medini et al., 2005; Tettelin et al., 2005; Hogg et al., 2007). The pan-genome consists of core genes, present in all sequenced strains, dispensable, or periphery, genes, present in a subset of strains, and unique, strain-specific genes. The pan-genome is said to be open if upon addition of new strains its size continues to grow, or closed, if at some point it saturates (Medini et al., 2005).

Fractions of the pan-genome may differ not only in size, but also in the functional composition (Gordienko, Kazanov, and Gelfand, 2013). In general, core genes encode housekeeping functions, while dispensable and unique genes confer selective advantages such as adaptation to particular niches, e.g. colonization of different hosts for pathogens, or antibiotic resistance (Muzzi, Masignani, and Rappuoli, 2007). So one may expect that genes from different fractions of the pan-genome evolve in different modes, including gene gain/loss rate, frequency of horizontal gene transfer, and selective pressure (Sarkar and Guttman, 2004; Wolf et al., 2016).

A consequence of the highly dynamic nature of bacterial genomes is frequent genomic rearrangements. Large inversions across the replication axis, deletions and insertions have been observed in *S. pneumoniae* (Camilli et al., 2011; Williams et al., 2012), *S. suis* (Yao et al., 2015; Athey et al., 2015), *S. pyogenes* (Hamada, Kawabata, and Nakagawa, 2015). The inversions have been suggested to rebalance the chromosomal architecture affected by insertions of large DNA segments (Camilli et al., 2011). The majority of these rearrangements occur at genome areas encoding transposases. Other genomic rearrangements occur at rRNA operons or sites encoding phage integrases and/or phage-related proteins.

The available genomes of *S. pneumoniae* strains have been classified into three categories according to their collinearity, and the obtained synteny types are largely consistent with the phylogenetic relationships seen among these strains (Zhang et al., 2011).

Genome arrangement may have profound effects on a bacterial phenotype. Rearrangements can disrupt genes, create new genes by fusion of gene parts, or change gene expression. One example of such inversions is truncation of the so-called *srtF* pilus island in *S. pneumoniae* NSUI060 (Athey et al., 2016). In *S. pyogenes* M23ND, genomic rearrangements resulted in re-clustering of a broad set of *CovRS*-regulated, actively transcribed genes, including virulence factors and metabolic genes, to the same leading strand. This may provide a potential advantage by creating spatial proximity to the transcription complexes, which may contain the global transcriptional regulator, *CovRS*, and RNA polymerases, in turn allowing for efficient transcription of the genes required for growth, virulence, and persistence (Bao et al., 2016).

Here we describe the pan-genomes of *S. pneumoniae*, *S. pyogenes*, and *S. suis*, their functional composition, the pattern of selection acting on genes and intergenic regions from different parts of the pan-genome, and genomic rearrangements within species.

## Methods

### Genome sequences

The selection of the species was based on the number of available strain genomes. We analyzed 25 strains of *Streptococcus suis*, 50 strains of *Streptococcus pyogenes*, and 28 strains of *Streptococcus pneumoniae*, all available complete genomes as of June 2016 (Table S1 and Fig S1). The complete genomes were downloaded from the NCBI Genome database (NCBI, 2017).

### Construction of orthologous groups (OGs)

We constructed orthologous groups using Proteinortho v5.13 with the default parameters (Lechner et al., 2011). Each gene was thus assigned to an orthologous group or labeled as a singleton.

### Estimation of the pan-genome size

We estimated the size of a pan-genome with the Chao algorithm from the micropan R-package (Snipen and Liland, 2015).

### Assignment of Gene Ontology (GO) terms to orthologous groups

To assign GO terms to genes, we used Interproscan (Jones et al., 2014). A GO term was assigned to an orthologous group, if it was assigned to at least 90% of genes in this group. To determine overrepresented functional categories, we used GOstat (Beißbarth and Speed, 2004). The fit by theoretical models was estimated using the Akaike information criterion (AIC) (Hurvich and Tsai, 1989).

### Assignment of KEGG Orthology (KO) categories to orthologous groups

Initially, we assigned KO categories to genes with GhostKOALA (Kanehisa, Sato, and Morishima, 2016). Then a KO category was assigned to an orthologous group, if it was assigned to at least 90% of genes in this group. KO terms were divided into supercategories “Genetic Information Processing”, “Metabolism”, “Cellular Processes”, “Environmental Information Processing”, and “other” based on the KEGG hierarchy classification.

### Prediction of virulence-related orthologous groups

We found virulence-related genes with MP3 (threshold 0.2) (Gupta et al., 2014) that combined a support vector machine classifier trained on virulence factors from MvirDB (Zhou et al., 2006) and a hidden Markov model classifier based on Pfam domains present in virulence factors. Orthologous group was considered virulence-related if at least 10% of its members were predicted to be virulence-related.

### Prediction of prophage insertions

To predict potential prophages, we used web server PHAST (Zhou et al., 2011).

### *pN/pS* calculation

To estimate the number of synonymous (*pS*) and non-synonymous (*pN*) polymorphisms, we aligned amino acid sequences of proteins using MUSCLE (Edgar, 2004) and then reconstituted the corresponding nucleotide alignment. Then we calculated *pN* and *pS* using the KaKs-Calculator Toolbox v2.0 with the Modified version of the Yang-Nielsen (MYN) method (Zhang et al., 2006). Multiple substitutions were accounted for using the Jukes-Cantor correction (Jukes, Cantor, and Munro, 1969). For these calculations, we considered different *Streptococcus* species separately. For each species and each orthologous group not containing paralogos, we performed pairwise comparisons of all strains and assigned the median *pN/pS* ratio to this group.

### Selection in intergenic regions

We extracted intergenic regions from .gbk files downloaded from the NCBI Genome database. We removed intergenic regions shorter then 50 bp. Out of the remaining intergenic regions we constructed the sample of upstream fragments in the following way. We extracted 100 bp upstream fragments for all intergenic regions longer than 100 bp (Gordon et al., 2005; Burden, Lin, and Zhang, 2004). For intergenic regions shorter than 100 bp its complete sequence was considered as an upstream fragment.

We estimated the number of positions under negative selection in two ways. Firstly, we simply calculated substitutions in upstream fragments. For that, we considered all pairs of strains from one species, extracted aligned upstream fragments of orthologous genes from the multiple genome alignment, and counted nucleotide substitutions with the Jukes-Cantor correction.

Secondly, we applied the method from (Tsoy et al., 2012) that calculates the fraction of positions under negative selection by comparing conservation statistics of multiple sequence alignments of orthologous upstream fragments from strains of two closely related species.

### Detection and analysis of large insertions/deletions (indels)

In orthologous upstream fragments we considered all indels of length at least 6 nucleotides, observed in at least two strains, and not located at the alignments termini (to reduce the bias from misalignment of fragment termini and varying length of upstream regions).

### Identification of candidate transcription-factor binding sites

We scanned for candidate binding sites in upstream fragments with FIMO (Grant, Bailey, and Noble, 2011), using positional weight matrices downloaded from PRODORIC (Münch et al., 2003). Candidate binding sites were filtered using the FDR correction for multiple testing (*q* < 0.05).

### Gene composition of the leading and lagging strands

We identified origin (*OriC*) and terminus (*Ter*) of replication analyzing GC-skew plots. Based on the *OriC* and *Ter* locations, we determined the strands for genes from different fractions of the pan-genome. To test the statistical significance of differences between the pangenome fractions, we performed a permutation test by shuffling genes between pan-genome fractions (retaining the fractions sizes) 250 times, thus obtaining the distribution of differences between the fractions under the random null model, and compared the observed differences with this distribution. Calculated differences with *p*-value satisfying the threshold with the Bonferroni correction for multiple testing were considered as statistically significant.

Statistical significance of over-representation of inter-replichore inversions was calculated as the probability of a given number of inter-replichore inversions in the set of inversions with given lengths. The probability of occurrence of the origin or the terminator of replication within the inversion was calculated as the ratio of the inversion length to the replichore length.

### Construction of phylogenetic trees

For construction of phylogenetic trees we used concatenated aligned amino acid sequences of all core genes reverse translated to nucleotide alignment. Then maximum likelihood trees were constructed by RAxML (Stamatakis, 2006) with default parameters.

### Synteny blocks and rearrangements history

Synteny blocks were constructed using the Sibelia algorithm (Minkin et al., 2013) with default parameters for whole-genome nucleotide alignments. Blocks observed in a genome more than once were filtered out. The history of inversions was reconstructed using the MGRA algorithm (Avdeyev et al., 2016).

### Detection of gene inflow

To detect genes horizontally transferred into species, we used the following model. If a gene with a mosaic phyletic pattern has been inherited vertically from the common ancestor and lost by several genomes, we expect to find it at the same syntenic region in the remaining strains. Genes not satisfying this condition are candidates for having been obtained horizontally. For this analysis, we excluded genes whose universal neighbours were affected by the reconstructed rearrangements, that is, genes located at or near boundaries of synteny blocks.

## Results and discussion

### Pan-genome and its fractions

We constructed 5742 OGs comprising 192782 genes (the number of genes in a genome assigned to OGs was 1872 *±* 178 with the median 1857, Tables S1 and S2). The number of singletons was 48 *±* 53, median=22.

For strains of each species and for combinations of species we performed the standard pan-genome analysis to characterize the distribution of OGs by strains and to estimate the sizes of core and pan-genomes. The distribution of OGs by strains had a typical U-shape (Hogg et al., 2007; Gordienko, Kazanov, and Gelfand, 2013; Koonin and Wolf, 2008) (Fig. 1A), that could be fitted by a sum of three exponents (as in (Koonin and Wolf, 2008)), describing the common core, periphery genes, and unique genes, or by a sum of two power law functions (as in (Gordienko, Kazanov, and Gelfand, 2013)), that divide the pan-genome into almost universal and almost unique genes (Fig. 1B). These two fits had almost equal R-squared values, but based on the AIC, the sum of three exponents was slightly more significant. When all three species were combined, the U-curve had additional minor peaks reflecting species-specific genes (Fig. 1A).

**Figure 1:**
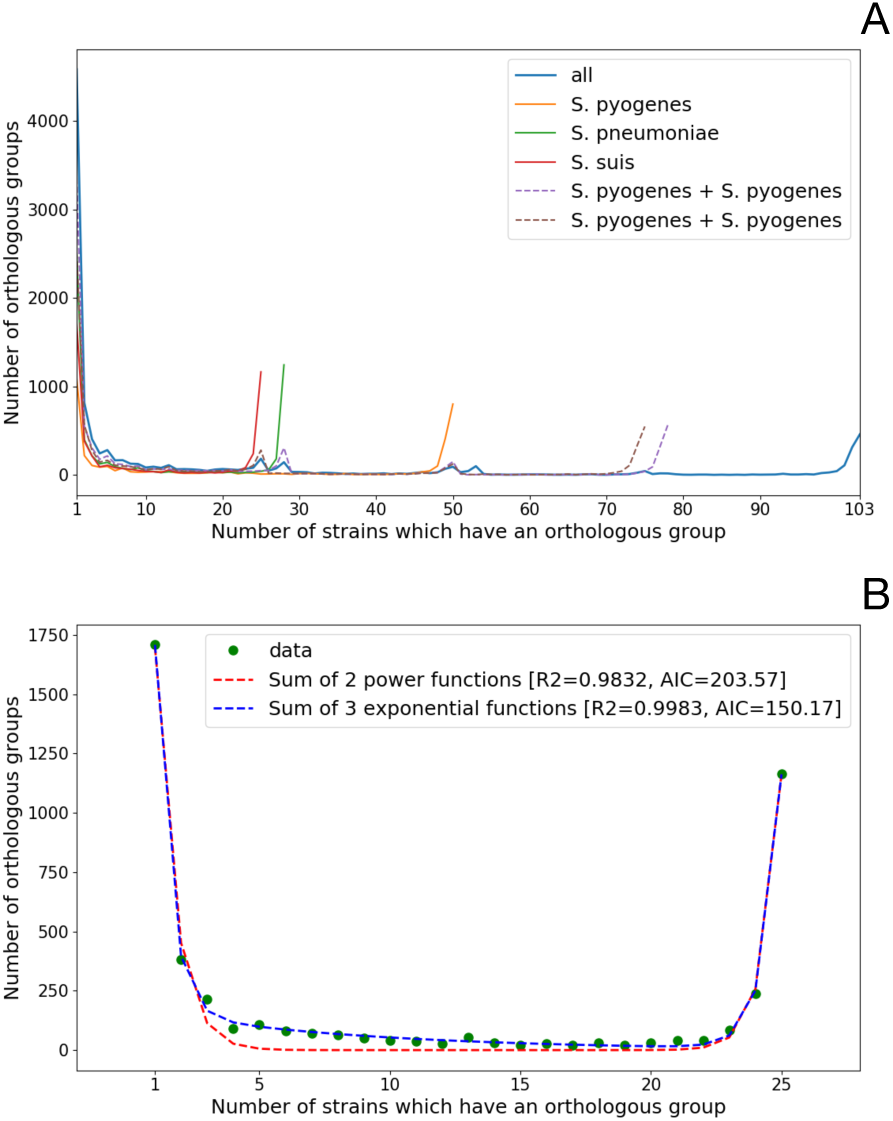
Distribution of orthologous groups (OGs) by the number of strains in which they are present. (A) For all analyzed strains (25 S. suis, 28 S. pneumoniae, and 50 S. pyogenes). (B) For 25 strains of S. suis with fitting by the sum of three exponential functions and by the sum of two power functions.

The core genome of the three species converged to 458 genes (core and pan-genomes of separate species described in Table 1). The pan-genome was open, exceeding 10300 genes (Fig. 2A). Species-specific pangenomes are also open, with core genomes accounting for approximately half of genes in any given genome. The Chao approximation of the total pan-genome size was 23217. The fraction of unique genes in a genome was less than 4%, with the highest fraction of unique genes in *S. pneumoniae*, and the lowest one in *S. pyogenes* (Fig. S2). The latter observation is partially explained by the presence of some very close strains in the analyzed genome set.

**Figure 2:**
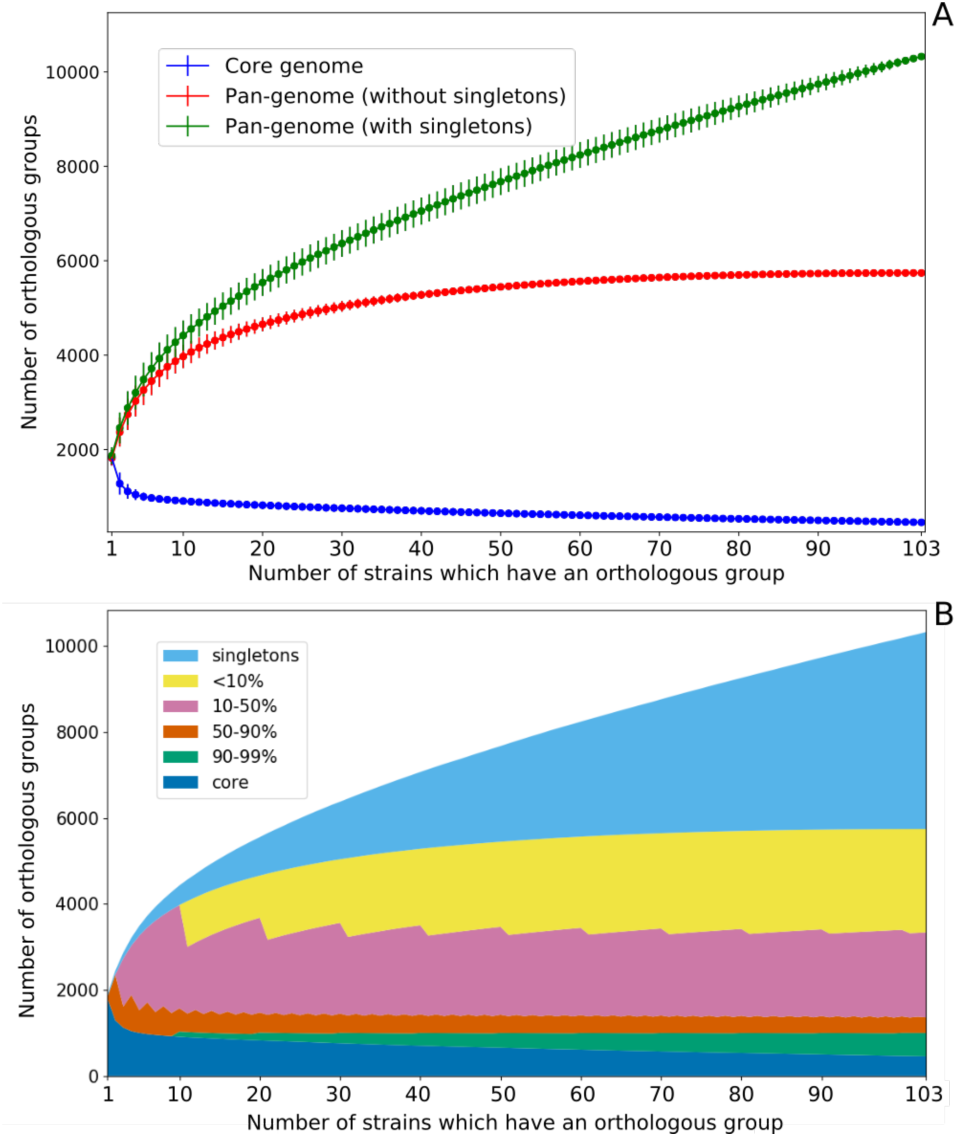
Sizes of the pan-genome fractions. (A) Sizes of the core genome, the pan-genome without singletons, and the total pan-genome with singletons as functions of the number of the analyzed strains. (B) Number of orthologous groups (OGs) are present in a given fraction of strains. Core OG present in all strains, singletons are present in only one strain each. All fractions reach saturation, while the pangenome continues to grow due to singletons.

**Table 1:** Sizes of pan-genome fractions.

| Species | Core genes | > 90% | > 50% | > 10% | Not species-specific | Total pan-genome | Chao estimation |
| --- | --- | --- | --- | --- | --- | --- | --- |
| All | 458 | 999 | 1361 | 3339 | 5742 | 10326 | 23217 |
| *S. suis* | 1164 | 1486 | 1803 | 2582 | 3264 | 4672 | 8562 |
| *S. pneumoniae* | 1243 | 1480 | 1838 | 2768 | 3410 | 5672 | 6477 |
| *S. pyogenes* | 800 | 1403 | 1588 | 2258 | 2843 | 3757 | 13680 |

As in (Gordienko, Kazanov, and Gelfand, 2013), we split the pan-genome into percentile fractions by considering OGs present in at least a given fraction of strains. All such pan-genome fractions reach saturation after addition of the first few strains, an exception being the core genome, that continues shrinking, although at a decreasing rate, and the total pan-genome that grows, mostly due to strain-specific, unique genes. If unique genes are excluded, the total pan-genome becomes closed and converges to about 5750 genes (Fig. 2B and Table 1).

We also considered the OG distribution in all three species simultaneously as a plot in three dimensions (Fig. 3). Excluding singletons, the largest group of OGs was formed by species-specific periphery (1136 in *S. pneumoniae*, 891 in *S. suis*, 922 in *S. pyogenes*), then OGs from the common core of the three species (458 OGs; or 825 OGs for a more relaxed definition with OG allowed to be absent in one strain in each species), then OGs belonging to the inner space of the plotted cube, i.e. to the common periphery of all three species (270 OGs), species-core OGs (114 in *S. pneumoniae*, 126 in *S. suis*, 87 in *S. pyogenes*), and, finally, some OGs formed common cores of species pairs to the exclusion of the third species (93 for *S. pneumoniae* and *S. suis*, 30 for *S. suis* and *S. pyogenes*, 12 for *S. pneumoniae* and *S. pyogenes*, reflecting closer phylogenetic relationships between *S. pneumoniae* and *S. suis*).

**Figure 3:**
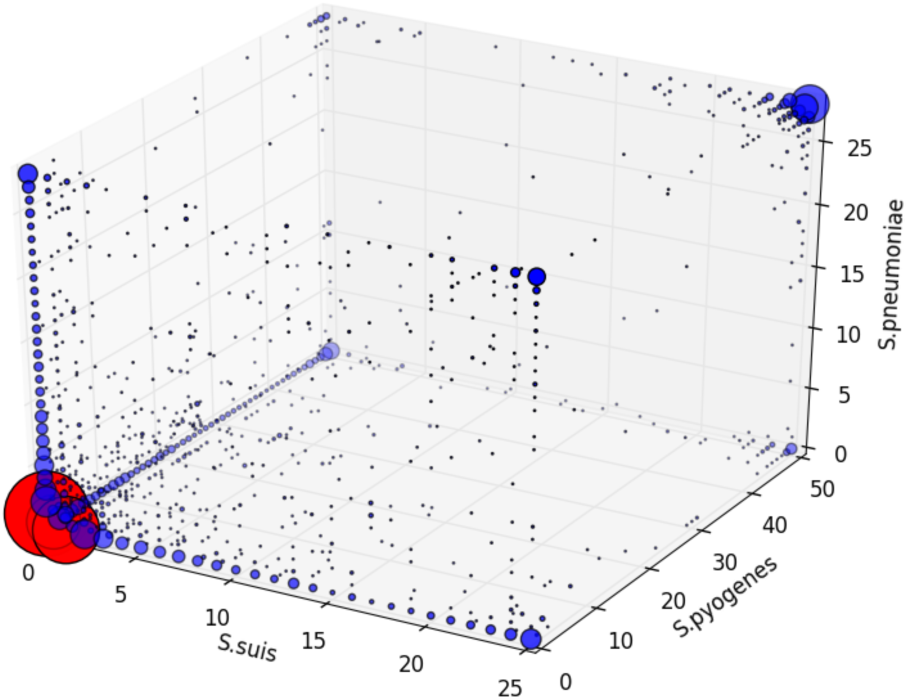
Distribution of orthologous groups (OGs) by the number of strains in which they are present as a plot in three dimensions. Axes correspond to species. Size of dots reflects the number of OGs. Red dots marks singletons OGs. Most dots reside at edges or in corners of the cube plot.

### Distribution of GO terms across pangenome fractions

Interproscan (Jones et al., 2014) provided at least one GO term to 127672 genes. These assignments are largely consistent, as members of an orthologous group tend to be assigned the same GO term (Fig. S3). Requiring that at least 90% of proteins from an OG share the GO term, we assigned GO terms to 2969 orthologous groups.

The distribution of orthologous groups with determined GO terms across the pan-genome, given in Fig. 4A, shows that core-genome groups tend to be more often assigned GO terms than genes from the unique fraction of the pan-genome. Indeed, the unique genes mainly had no GO terms (Fig. 4A, “Strain-specific OGs”) or KEGG KO terms (Fig. 4B, “Strain-specific OGs”, Fig. S4) and were annotated as “hypothetical proteins”, hence likely encoding mobile elements and phage-related proteins or simply resulting from genome misannotation (Fig. S5). However, some important gene groups also fell in this category, as 15% of unique genes were predicted to be virulence-related (Fig. S6). The exact fraction of functionally relevant genes in this group is hard to estimate, as the absence of homologs makes functional annotation almost impossible (although gene calling artifacts in some cases may be recognized by the comparison of strains).

**Figure 4:**
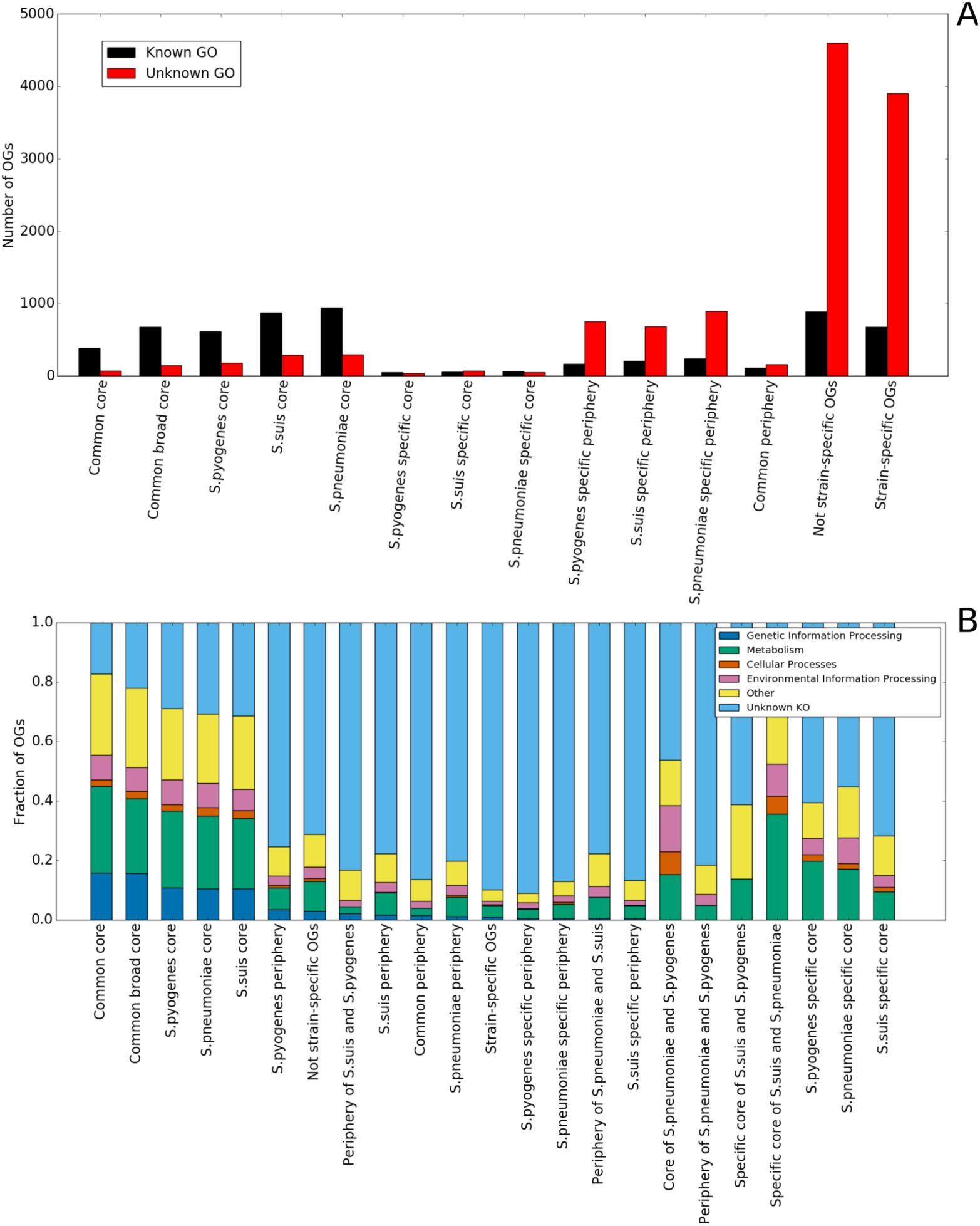
Distributions of orthologous groups (OGs). (A) with or without a GO term and (B) with known or unknown high-level KEEG KO category across the pan-genome fractions. Pan-genome fractions: Common core - OGs present in all strains, Common broad core - OGs missing at most in one strain of each species, S. pyogenes core - OGs present in all S. pyogenes strains, S. suis core - OGs present in all S. suis strains, S. pneumoniae core - OGs present in all S. pneumoniae strains, S. pyogenes specific core - OGs present in all S. pyogenes strains and absent in other species, S. suis specific core - OGs present in all S. suis strains and absent in other species, S. pneumoniae specific core - OGs present in all S. pneumoniae strains and absent in other species, S. pyogenes specific periphery - OGs present in some S. pyogenes strains and absent in other species, S. suis specific periphery - OGs present in some S. suis strains and absent in other species, S. pneumoniae specific periphery - OGs present in some S. pneumoniae strains and absent in other species, common periphery - OGs present in some but not all strains of each species, not strain-specific OGs - all OGs excluding singleton OGs, strain-specific OGs - singleton OGs, S. pyogenes periphery - OGs present in some S. pyogenes strains, periphery of S. suis and S. pyogenes - OGs present in some S. suis and S. pyogenes strains, S. suis periphery OGs present in some S. suis strains, S. pneumoniae periphery - OGs present in some S. pneumoniae strains, periphery of S. pneumoniae and S. suis - OGs present in some S. pneumoniae and S. suis strains, specific core of S. pneumoniae and S. pyogenes - OGs present in all S. pneumoniae and S. pyogenes strains and absent in S. suis strains, periphery of S. pneumoniae and S. pyogenes - OGs present in some S. pneumoniae and S. pyogenes strains, specific core of S. suis and S. pneumoniae - OGs present in all S. suis and S. pneumoniae strains and absent in S. pyogenes strains, specific core of S. suis and S. pyogenes - OGs present in all S. suis and S. pyogenes strains and absent in S. pneumoniae strains.

Overrepresented functional categories in different fractions of the pan-genome with regards to the described cube representation are shown in Table S2. The common core genome and weakly species-specific cores, that is genes observed in all strains of one species and some strains of the remaining species, are enriched with genetic information processing GOs, such as translation, ribosome, gene expression, RNA, and all kinds of metabolic processes. The periphery is enriched in a small set of functions, including response to other organisms and pathogenesis (this fraction features the highest percent of predicted virulence-related genes, Fig. S6), in particular, sialidase activity (*S. pyogenes*, *S. pneumonie*), DNA binding and some carbohydrate-related functions (*S. pneumoniae*, *S. suis*), as well as transcription factors (*S. pyogenes*). Strain-specific genes are mainly enriched in transposase activity, DNA recombination and DNA integration, consistent with mobile elements origin of strain-specific genes (Wolf et al., 2016); in addition, these categories are enriched in orthologous groups from the common periphery, that is, among genes present in a fraction of strains from all three species. Species-specific cores are enriched in vitamin biosynthesis (*S. pneumonie*), transport, histidine and lactose metabolism, and response to oxidative stress (*S. pyogenes*), and iron transport, amino acid metabolic processes, and regulation of transcription (*S. suis*).

The distribution of KEEG KO categories across the pan-genome is shown in Fig. 4B. The fraction of orthologous groups assigned with a KO category decreases when moving from the core genome to the periphery and then to strain-specific genes. Most orthologous groups related to “Genetic Information Processing”, that can be considered as most essential groups, correspond to the common core, followed by the periphery and then strain-specific genes; no such orthologous groups were found among species-specific cores.

Hence, the functional distribution agrees with the pan-genome model in which the core is responsible for information and most metabolic processes, the periphery performs fine-tuning of bacteria to specific ecological niches, and strain-specific genome fraction is comprised mainly of mobile elements-related genes (Carlos Guimaraes et al., 2015).

Genes from the common core show a weak preference to the leading strand, whereas periphery and strain-specific genes tend to be located at the lagging strand (Table 2). The leading strand preference of the core genes may be associated with their higher transcription level and/or with essentiality of these genes (Zheng et al., 2015). However, this difference is not very strong, as *Streptococcus* feature a general, strong bias with about 80% of genes located at the leading strand.

**Table 2:** Strand preference of genes from pan-genome fractions. Here, periphery is defined as genes present in some strains of a given species and absent in other species. * - statistically significant.

| Pan-genome fraction (number of OGs) | Percent of genes on leading strand | Standart deviation |
| --- | --- | --- |
| All genes (5742 OGs + 4584 singletons) | 79.6 | 1.4 |
| Non strain-specific OGs (5742) | 79.8 * | 1.4 |
| Common core (458) | 83.9 * | 0.6 |
| *S.pneumoniae* specific core (114) | 79.6 | 0.9 |
| *S.pyogenes* specific core (87) | 73.5 * | 0.2 |
| *S.suis* specific core (126) | 78.8 | 1.6 |
| *S.pneumoniae* periphery (1136) | 75.2 * | 5.2 |
| *S.pyogenes* periphery (922) | 75.5 * | 3.6 |
| *S.suis* periphery (891) | 78.3 * | 3.3 |
| Common periphery (270) | 77.8 * | 6 |
| Strain-specific OGs (4584) | 70.1 * | 15.2 |

### Selection regime in the pan-genome fractions

Genes from the core genome encoding essential, housekeeping functions should evolve under higher purifying selection (Schloissnig et al., 2013) yielding lower *pN/pS* ratio, compared to dispensable genes from the periphery genome. Indeed, as shown in Fig 5A and Fig. S7A, the *pN/pS* ratio is the smallest for the core genome (Mann–Whitney, *p*<0.01 for comparisons of core and periphery fractions within the same species).

**Figure 5:**
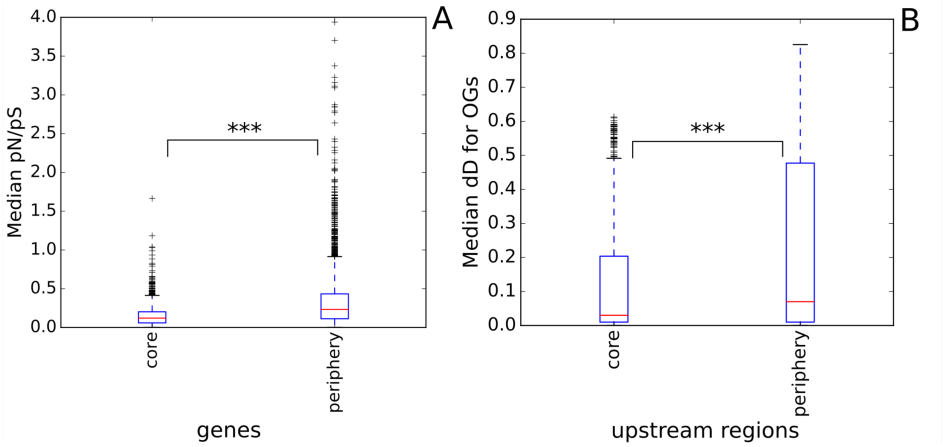
Selection regime in the pan-genome fractions. (A) Median value of the pN/pS ratio with the Jukes-Cantor correction for genes and (B) median number of nucleotide substitutions in upstream regions (dD). Significance level is indicated with stars (p-value < 0.001).

### Selection in gene upstream regions and transcriptional regulation

In addition to protein-coding genes, purifying selection acts on regulatory elements in intergenic regions. In this and the next sections, we attempt to quantify this selection by determining the fraction of intergenic nucleotide positions evolving under negative selection and by comparing regions that are deleted in some strains with universal intergenic regions.

The median fraction of nucleotide substitutions (with the Jukes-Cantor correction) in intra-species alignments of orthologous upstream regions was 5.6%. The distribution of the number of nucleotide substitutions with the Jukes-Cantor correction, *dD*, is shown in Fig. 5B and Fig. S7B. The fraction of the pan-genome with the lowest number of substitutions is the core genome. Hence, not only the core genes, but their expression level and regulation are likely to be conserved.

In inter-species alignments, conserved columns may indicate functional conservation or simply insufficient time after speciation to accumulate mutations in all non-essential positions. To estimate the number of hidden non-conserved positions we used the method from (Tsoy et al., 2012). We have calculated that only 10-20% of positions in the upstream regions evolve under purifying selection (Fig. S8). However, this may be an underestimate due to the large distance between the analysed species and the low number of conserved positions.

### Inserted and deleted fragments in intergenic regions are not neutral

Many alignments of orthologous upstream regions contained extended insertions and/or deletions (indels). One might expect that the sequences within indels are selectively neutral. However, the indel fragments demonstrate strong sequence conservation (in the remaining genomes), and the level of conservation increases with the indel length (Fig. 6, Fig. S9). One possible explanation could be horizontal transfer of regulatory sequences enabling fast change of the level and mode of expression for the adjacent gene(s) (Oren et al., 2014). However, computational scanning for candidate binding sites of transcription factors [reference to the database / search tool] in the indel fragments has not produced an excess of candidate sites compared with control, random fragments from the same upstream regions, controlled for length. This might be due to low recall of the recognition rules, noise in predictions, and the fact that evolving intergenic regions may contain genes for regulatory RNAs (Čuklina et al., 2016; Smirnov et al., 2017).

**Figure 6:**
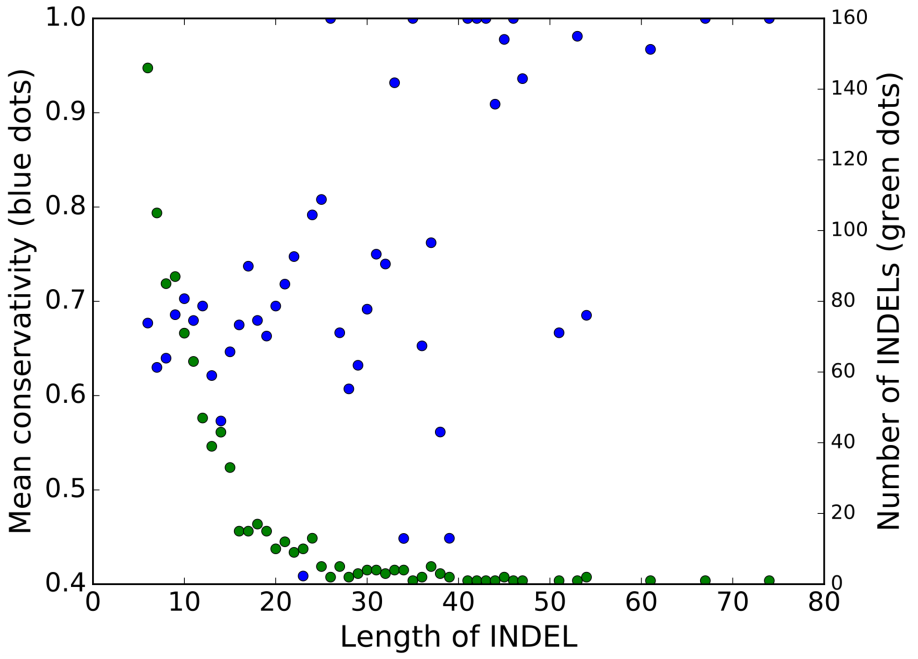
Dependence of the mean conservation level of nucleotides in indels in upstream regions in genome of S. pyogenes on the indel size. Blue dots correspond to the mean conservation level, green dots correspond to the number of indels of this size.

### Genomic rearrangements

An important mode of genome evolution is rearrangements of chromosome fragments. In prokaryotes with single chromosomes the prevalent type of rearrangements are the symmetrical inversions around the origin of replication (Bochkareva et al., 2018; Cossu et al., 2017; Repar et al., 2017; Wang et al., 2017). While a number of inversions in some *Streptococcus* strains have been described (Zhang et al., 2011), the increased phylogenetic coverage allowed us to actually map the events to the phylogenetic tree.

Synteny blocks were obtained using whole-genome alignments for each species. Only blocks present in all strains were used for the reconstruction of inversions. As a result, 13 inversions for *S. pneumoniae*, 21 inversions for *S. suis*, and 26 inversions for *S. pyogenes* were identified. Mapping these events to phylogenetic trees (Fig. S10) revealed cases of parallel inversions in all three species.

To check whether these parallel events were caused by homologous recombination, we constructed trees based on genes involved in these events (Fig. S11). In all cases, strains with parallel inversion did not cluster together, as would be expected in the case of spread by homologous recombination involving these regions.

Previously, inversions in *Streptococcus* spp. were explained by selection to rebalance the replichore architecture affected by insertion of prophages (Camilli et al., 2011). To check this hypothesis, we compared lengths of prophage regions in strains that contained the same inversion, and vice versa the number of inversions in strains with the same rate of prophage insertions (Table S1). No correlation between the rates of prophage insertions and inversions was observed.

All inversions were bounded by mobile elements or clusters of rRNA except one event in the *S. pneumoniae* subtree. This inversion of length 15 kB was found in four separated branches and breakpoints were formed by genes encoding the surface antigen proteins PhtB and PhtD from a family characterized by the presence of several histidine triad (HxxHxH) motifs. PhtD and PhtB are relatively large proteins with lengths about 850 amino acids thought to be involved in multiple functions, including metal ion homeostasis, evasion of complement deposition, and adherence of bacteria to host cells (Plumptre, Ogunniyi, and Paton, 2012). In pairs of strains with and without the inversion, these proteins are composed of two independent parts with the inversion breakpoint in the middle of the gene (Fig. 7).

**Figure 7:**
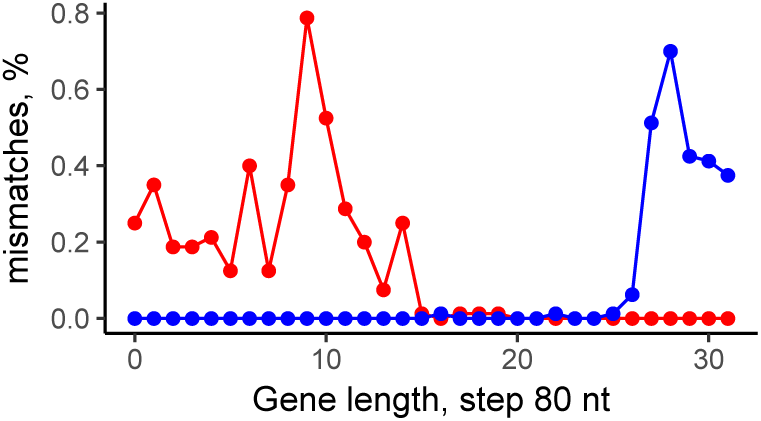
Number of mismatches in alignments of the histidine-triad proteins in S. pneumoniae ST556 and S. pneumoniae TCH8431/19A. The red points and blue points show the local dissimilarity level of PhtB from S. pneumoniae ST556 vs PhtB in S. pneumoniae TCH8431/19A and PhtD from S. pneumoniae TCH8431/19A, respectively.

As phase variation is known to be important for *S. pneumoniae* infection (Li et al., 2016), the observed parallel inversion might indicate the action of an antigen variation mechanisms, which is important, as PhtD is a candidate for a next-generation pneumococcal vaccine (Yun et al., 2015).

As more than 80% core genes in *Streptococcus* spp. are found on the leading strand (see above), one would expect strong selection against intra-replichore inversions, as they do switch genes between leading/lading strands. Indeed, inter-replichore inversions are over-represented (57 events of 62, the *p*-value = 9 × 10^−14^) (Fig. 8).

**Figure 8:**
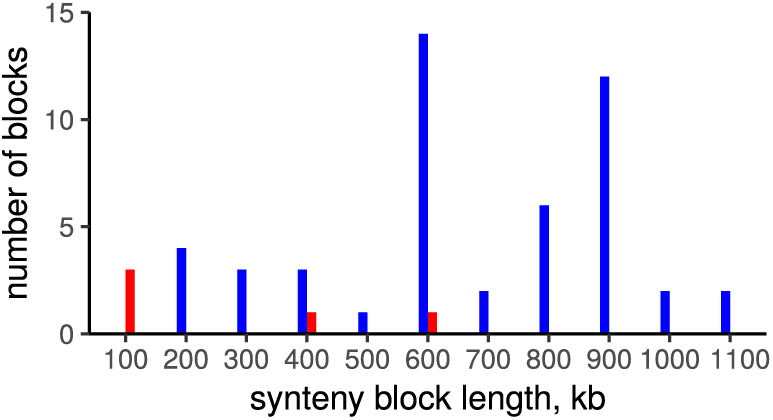
Distribution of inversion lengths. Red color corresponds to intra-replichore inversions; blue color, inter-replichore inversions.

### Detection of gene inflow

To identify genes horizontally transferred after the divergence of the studied *Streptococcus* spp. and further spreading between the strains, we selected genes that were not unique and that were not common for at least one *Streptococcus* species (referred to periphery genes). Positions of single-copy, universal genes were analyzed to construct syntenic regions for all strains and to compile a set of periphery genes occurring in different syntenic regions and, therefore, likely being spread by horizontal gene transfer. The set comprised 277 orthologous groups that is about 7% of single-copy periphery genes. More than a half of these genes are rare in all three *Streptococcus* species; others are rare in at least one species (Fig. 9).

**Figure 9:**
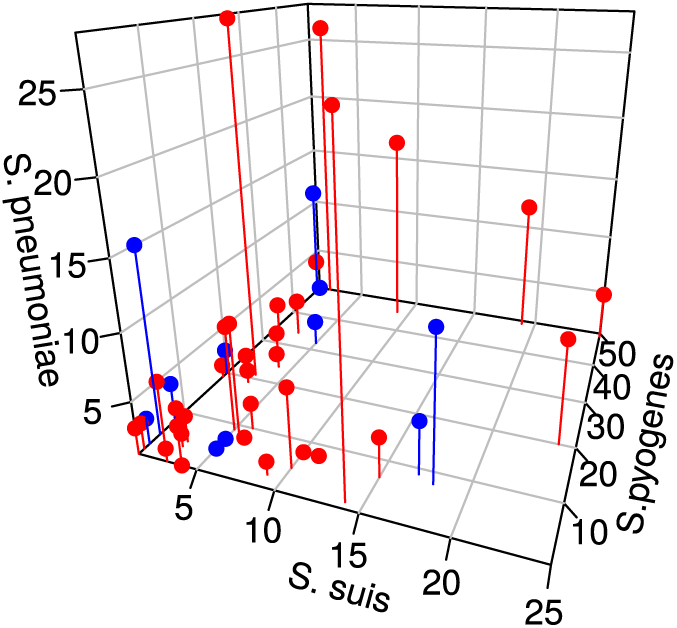
Distribution of genes with non-conserved genome positions. One point corresponds to one OG. Blue points correspond to OGs that have consistent trees; red points, OGs that have inconsistent trees.

To confirm horizontal gene transfer, we constructed phylogenetic gene trees for all orthologous groups that were present at least in two *Streptococcus* species and at least in two strains in each species and checked whether each species were monophyletic, that is, formed a separate branch in these trees (Table 3). Most groups (88%) yielded trees with monophyletic species (consistent trees) and had conserved location in all genomes, indicating vertical inheritance from a common ancestor. About a half of groups with inconsistent trees had conserved genome positions that may be explained by homologous recombination; the remaining half had non-conserved location. The set of 48 orthologous groups with inconsistent trees and non-conserved positions are candidates for horizontal transfer between species.

**Table 3:** Statistics of periphery genes.

|  | Conserved location | Non-conserved location | All |
| --- | --- | --- | --- |
| Consistent tree | 1181 | 128 | 1309 |
| Inconsistent tree | 44 | 48 | 92 |
| All | 1225 | 176 | 1401 |

An analysis of overrepresented GO terms in the set of orthologous groups with non-conserved positions, compared with all non-core groups (Table 4), yielded many functions involved in interaction with DNA (DNA binding, nucleic acid binding, sequence-specific DNA binding), that may be linked to regulation, but also to mobile elements (DNA integration). Other overrepresented functions such as ATP hydrolysis coupled proton transport, energy coupled proton transport against electrochemical gradient, proton-transporting V-type ATPase complex, etc., all are likely linked to the V-type ATPase, that is thought to be horizontally transfered from archaea (“Inventing the dynamo machine: the evolution of the F-type and V-type ATPases”).

**Table 4:** Overrepresented GO terms in genes with non-conserved location, compared with all non-core genes.

| GO term | Genes with non-conserved location out of all genes | P-value | Description |
| --- | --- | --- | --- |
| GO:0003677 | 51 out of 603 | $1.5 * 10^{-3}$ | DNA binding |
| GO:0015991 | 4 out of 5 | $2.2 * 10^{-3}$ | ATP hydrolysis coupled proton transport |
| GO:0015988 | 4 out of 5 | $2.2 * 10^{-3}$ | energy coupled proton transport against electrochemical gradient |
| GO:0003676 | 58 out of 762 | $3.3 * 10^{-3}$ | nucleic acid binding |
| GO:0016469 | 4 out of 6 | $3.8 * 10^{-3}$ | proton-transporting two-sector ATPase complex |
| GO:0006818 | 4 out of 7 | $6.2 * 10^{-3}$ | hydrogen transport |
| GO:0015992 | 4 out of 7 | $6.2 * 10^{-3}$ | proton transport |
| GO:0043565 | 14 out of 98 | $6.8 * 10^{-3}$ | sequence-specific DNA binding |
| GO:0015074 | 13 out of 100 | $2.6 * 10^{-2}$ | DNA integration |
| GO:0033176 | 2 out of 2 | $5.5 * 10^{-2}$ | proton-transporting V-type ATPase complex |
| GO:0033179 | 2 out of 2 | $5.5 * 10^{-2}$ | proton-transporting V-type ATPase, $V_0$ domain |

### Conclusions

In a typical *Streptococcus* genome, one quarter of genes belong to the genus core genome; one quarter, to the species-specific core; most other genes are periphery ones, and a minority are strain-specific. The core genome is enriched with information-process and main metabolic functions and depleted with mobile elements and phage-related genes; the periphery fraction is enriched with niche-specific metabolic functions, including pathogenesis-related ones; and strain-specific genes are enriched with hypothetical genes and mobile elements, but also contain many virulence-related genes. The core genes demonstrate a lower level of substitutions and show a stronger tendency to be located on the leading strand.

Upstream regions of the core genes have fewer substitutions than average, which might reflect a stronger conservation of their regulation or more complex regulation, yielding a larger density of transcription-factors binding sites and other regulatory structures. On the other hand, fragments of intergenic regions, that are deleted (or inserted) in some strains, are not less conserved than the surrounding regions, which might be a sign of newly evolving regulatory interactions.

The pan-genome of *Streptococcus* is open mainly due to strain-specific genes. The stable core genome performing housekeeping functions, broad periphery genome conferring niche adaptation, and a huge repertoire of strain-specific genes, potentially available to the genus members, provide the *streptococci* with a high level of adaptability. This emphasizes the importance of pan-genome studies of medically relevant bacteria, as their pathogenicity may be affected by rare periphery or even strain-specific genes.

The plasticity of *Streptococcus* genomes is due to horizontal gene transfer facilitated by the competence system (PA and PH, 2015). Indeed, the early proof that DNA carries genetic information was provided by experiments with *pneumococcus* (Griffith, 1928; Avery, MacLeod, and McCarty, 1944). At that, about 7% single-copy periphery genes occur in different syntenic regions. More than a half of these genes are rare in all *Streptococcus* species; others are rare in at least one species. The genes with inconsistent trees and nonconserved genome position have likely experienced horizontal transfer between species.

Another mechanism that leads to phenotype diversification is phase variation via intra-genomic recombination. The observed parallel inversion between paralogous genes encoding surface antigen protein in *S. pneumoniae* might indicate the action of an antigen variation mechanisms, which is important, as this protein is a candidate for a next-generation *pneumococcal* vaccine (Yun et al., 2015).

## Declarations

### Availability of data and materials

All sequences analyzed in this study were taken from GenBank. Accession numbers and details are available in Additional Table S1. Orthologous groups composition described in Additional Table S3 and GO term annotations are available in Additional Table S2.

### Author’s contributions

PVS, OOB and MSG conceived and designed the study; PVS, OOB and AAK analyzed the data; PVS, OOB and MSG wrote the paper. All authors read and approved the final version of the manuscript.

### Ethics approval and consent to participate

Not applicable

### Consent for publication

Not applicable

#### Acknowledgements

We are grateful to Marat Kazanov for sharing preliminary data.

### Additional Files

Additional file Fig. S1 — Phylogenetic tree of analyzed *Streptococcus* strains based on alignments of universal single-copied genes.

Additional file Fig. S2 — Distribution of the number of singletons in strains belong to different species.

Additional file Fig. S3 — Dependence of the number of orthologous groups (OGs) assigned a GO term on the threshold for GO term assignment. The threshold is the minimal fraction of genes from an ortologous group that have a GO term. Singleton are not considered.

Additional file Fig. S4 — Distribution of high-level KEEG KO categories across pan-genome fractions. (A) absolute values, (B) relative values of four major KO categories. Pan-genome fractions are defined as in Fig. 4.

Additional file Fig. S5 — Proportion of orthologous groups with hypothetical or mobile/phage related genes in (A) strain-specific OGs and in (B) not strain-specific OGs.

Additional file Fig. S6 — Distribution of orthologous groups with virulence-related genes across pangenome fractions. Number of OGs in each pan-genome fraction is shown in brackets. Pan-genome fractions are defined as in Fig. 4.

Additional file Fig. S7 — Distributions of (A) the median value of the *pN/pS* ratio with the Jukes-Cantor correction for genes and (B) median number of nucleotide substitutions in upstream regions (*dD*) of genes from OGs from different pan-genome fractions. The number of analyzed OGs from each pan-genome fraction is shown in brackets. The pan-genome fractions are defined as in Fig. 4.

Additional file Fig. S8 — Fractions of nucleotides under purifying selection in upstream fragments of core OGs as a function of the number of compared strains in pair-wise analysis of species.

Additional file Fig. S9 — Dependence of the mean conservation level of nucleotides in indels on the indel size. Blue dots correspond to the mean conservation level, green dots correspond to the number of indels of this size. (A) *S. pneumoniae*, (B) *S. suis*, and (C) *S. pyogenes*.

Additional file Fig. S10 — Phylogenetic trees based on genes involved in parallel inversions.

Additional file Fig. S11 — Phylogenetic trees for (A) *S. pneumoniae*, (B) *S. suis*, (C) *S. pyogenes* based on the alignments of universal single-copy genes. The numbers at tree branches show the numbers of inversions. Strains with parallel inversions are marked by color labels. Strains with the same inversion are marked by the same color.

Additional file Table S1 — List of analyzed *Streptococcus* strains.

Additional file Table S2 — Overrepresented functional categories in different fractions of the pangenome.

Additional file Table S3 — Orthologous groups compositions.

## Bibliography

Athey, TB et al. (2015). “Complex population structure and virulence differences among serotype 2 *Streptococcus suis* strains belonging to sequence type 28”. In: PLoS One 10.9, e0137760.

Athey, TB et al. (2016). “Population structure and antimicrobial resistance profiles of *Streptococcus suis* serotype 2 sequence type 25 strains”. In: PLoS One 11.3, e0150908.

Avdeyev, P et al. (2016). “Reconstruction of ancestral genomes in presence of gene gain and loss”. In: Journal of Computational Biology 23.3, pp. 150–164.

Avery, OT., CM. MacLeod, and M McCarty (1944). “Studies on the chemical nature of the substance inducing transformation of *Pneumococcal* types: induction of transformation by a desoxyribonucleic acid fraction isolated from *Pneumococcus* type III”. In: J. Exp. Med. 79.2, 137–158.

Bao, YJ et al. (2016). “Novel genomic rearrangements mediated by multiple genetic elements in *Streptococcus pyogenes* M23ND confer potential for evolutionary persistence”. In: Microbiology 162.8, pp. 1346–1359.

Beißbarth, T and TP Speed (2004). “GOstat: find statistically overrepresented Gene Ontologies within a group of genes”. In: Bioinformatics 20.9, pp. 1464–1465.

Bochkareva, OO et al. (2018). “Genome rearrangements and phylogeny reconstruction in textitYersinia pestis”. In: PeerJ 6, e4545.

Brown, JS, SM Gilliland, and DW Holden (2001). “A *Streptococcus pneumoniae* pathogenicity island encoding an ABC transporter involved in iron uptake and virulence”. In: Molecular microbiology 40.3, pp. 572–585.

Burden, S., Y-X Lin, and R Zhang (2004). “Improving promoter prediction Improving promoter prediction for the NNPP2.2 algorithm: a case study using *Escherichia coli* DNA sequences”. In: Bioinformatics 21.5, pp. 601–607.

Camilli, R et al. (2011). “Complete genome sequence of a serotype 11A, ST62 *Streptococcus pneumoniae* invasive isolate”. In: BMC Microbiol 11.25.

Carlos Guimaraes, L et al. (2015). “Inside the pangenome-methods and software overview”. In: Current genomics 16.4, pp. 245–252.

Cossu, M et al. (2017). “Flipping chromosomes in deep-sea archaea”. In: PLoS genetics 13.6, e1006847.

Čuklina, J et al. (2016). “Genome-wide transcription start site mapping of *Bradyrhizobium japonicum* grown free-living or in symbiosis–a rich resource to identify new transcripts, proteins and to study gene regulation”. In: BMC genomics 17.1, p. 302.

Edgar, RC (2004). “MUSCLE: multiple sequence alignment with high accuracy and high throughput”. In: Nucleic acids research 32.5, pp. 1792–1797.

Ferretti, JJ, DL Stevens, and VA Fischetti (2016). “Poststreptococcal autoimmune sequelae: rheumatic fever and beyond–*Streptococcus pyogenes*: basic biology to clinical manifestations”. In:

Gordienko, EN, MD Kazanov, and MS Gelfand (2013). “Evolution of pan-genomes of *Escherichia coli, Shigella* spp., and *Salmonella enterica*”. In: Journal of bacteriology 195.12, pp. 2786–2792.

Gordon, JJ et al. (2005). “Improved prediction of bacterial transcription start sites”. In: Bioinformatics 22.2, pp. 142–148.

Gottschalk, M and M Segura (2000). “The pathogenesis of the meningitis caused by *Streptococcus suis* : the unresolved questions”. In: Veterinary microbiology 76.3, pp. 259–272.

Grant, CE, TL Bailey, and WS Noble (2011). “FIMO: scanning for occurrences of a given motif”. In: Bioinformatics 27.7, pp. 1017–1018.

Griffith, F (1928). “The significance of *Pneumococcal* types”. In: Journal of Hygiene. Cambridge University Press. 27.2, 113–159.

Gupta, A et al. (2014). “MP3: a software tool for the prediction of pathogenic proteins in genomic and metagenomic data”. In: PloS one 9.4, e93907.

Hamada, S., S Kawabata, and I Nakagawa (2015). “Molecular and genomic characterization of pathogenic traits of group A *Streptococcus pyogenes*”. In: Proc Jpn Acad Ser B Phys Biol Sci 91.10, pp. 539–559.

Hogg, JS et al. (2007). “Characterization and modeling of the *Haemophilus influenzae* core and supragenomes based on the complete genomic sequences of Rd and 12 clinical nontypeable strains”. In: Genome biology 8.6, R103.

Hurvich, CM and CL Tsai (1989). “Regression and time series model selection in small samples”. In: Biometrika 76, pp. 297–307.

Jones, P et al. (2014). “InterProScan 5: genome-scale protein function classification”. In: Bioinformatics 30.9, pp. 1236–1240.

Jukes, TH, CR Cantor, HN Munro, et al. (1969). “Evolution of protein molecules”. In: Mammalian protein metabolism 3.21, p. 132.

Kanehisa, M, Y Sato, and K Morishima (2016). “BlastKOALA and GhostKOALA: KEGG tools for functional characterization of genome and metagenome sequences”. In: Journal of molecular biology 428.4, pp. 726–731.

Koonin, EV and YI Wolf (2008). “Genomics of bacteria and archaea: the emerging dynamic view of the prokaryotic world”. In: Nucleic acids research 36.21, pp. 6688–6719.

Krzyściak, W et al. (2013). “The pathogenicity of the *Streptococcus* genus”. In: European Journal of Clinical Microbiology and Infectious Diseases 32.11, pp. 1361–1376.

Lechner, M et al. (2011). “Proteinortho: detection of (co-) orthologs in large-scale analysis”. In: BMC bioinformatics 12.1, p. 124.

Li, J et al. (2016). “Epigenetic switch driven by DNA inversions dictates phase variation in *Streptococcus pneumoniae*”. In: PLoS Pathog. 12.7, e1005762.

Medini, D et al. (2005). “The microbial pan-genome”. In: Curr Opin Genet Dev 15.6, pp. 589–594.

Minkin, I et al. (2013). “Sibelia: A scalable and comprehensive synteny block generation tool for closely related microbial genomes”. In: 13th Workshop on Algorithms in Bioinformatics (WABI2013).

Mulkidjanian, AY et al. “Inventing the dynamo machine: the evolution of the F-type and V-type ATPases”. In: Nature reviews. Microbiology 5.11, p. 892.

Mullen, S (2015). “Review of pediatric autoimmune neuropsychiatric disorder associated with streptococcal infections”. In: Mental Health Clinician 5.4, pp. 184–188.

Münch, R et al. (2003). “PRODORIC: prokaryotic database of gene regulation”. In: Nucleic acids research 31.1, pp. 266–269.

Muzzi, A, V Masignani, and R Rappuoli (2007). “The pan-genome: towards a knowledge-based discovery of novel targets for vaccines and antibacterials”. In: Drug discovery today 12.11, pp. 429–439.

NCBI, Resource Coordinators (2017). “Database resources of the national center for biotechnology information.” In: Nucleic acids research 45.D1, p. D12.

Oren, Y et al. (2014). “Transfer of noncoding DNA drives regulatory rewiring in bacteria”. In: Proceedings of the National Academy of Sciences 111.45, pp. 16112–16117.

PA, Cheryl and William PH (2015). “Mechanisms of genome evolution of *Streptococcus*”. In: Infect Genet Evol 33, 334–342.

Plumptre, CD, AD Ogunniyi, and JC Paton (2012). “Polyhistidine triad proteins of pathogenic *streptococci*”. In: Trends Microbiol 20.10, pp. 485–493.

Repar, J et al. (2017). “Elevated rate of genome rearrangements in radiation-resistant bacteria”. In: Genetics 205.4, pp. 1677–1689.

Richards, VP et al. (2014). “Phylogenomics and the dynamic genome evolution of the genus *Streptococcus*”. In: Genome biology and evolution 6.4, pp. 741–753.

Sarkar, SF and DS Guttman (2004). “Evolution of the core genome of *Pseudomonas syringae*, a highly clonal, endemic plant pathogen”. In: Applied and Environmental Microbiology 70.4, pp. 1999–2012.

Schloissnig, S et al. (2013). “Genomic variation landscape of the human gut microbiome”. In: Nature 493.7430, p. 45.

Smirnov, A et al. (2017). “Discovery of new RNA classes and global RNA-binding proteins”. In: Current opinion in microbiology 39, pp. 152–160.

Snipen, L and KH Liland (2015). “Micropan: an R-package for microbial pan-genomics”. In: BMC bioinformatics 16.1, p. 79.

Stamatakis, A (2006). “RAxML-VI-HPC: maximum likelihood-based phylogenetic analyses with thousands of taxa and mixed models”. In: Bioinformatics 22.21, pp. 2688–2690.

Tettelin, H et al. (2005). “Genome analysis of multiple pathogenic isolates of *Streptococcus* agalactiae: implications for the microbial “pan-genome””. In: Proceedings of the National Academy of Sciences of the United States of America 102.39, pp. 13950–13955.

Tsoy, OV et al. (2012). “Evolution of transcriptional regulation in closely related bacteria”. In: BMC evolutionary biology 12.1, p. 200.

Wang, D et al. (2017). “Core-genome scaffold comparison reveals the prevalence that inversion events are associated with pairs of inverted repeats”. In: BMC genomics 18.1, p. 268.

Williams, TM et al. (2012). “Genome analysis of a highly virulent serotype 1 strain of *Streptococcus pneumoniae* from West Africa”. In: PLoS One 7.10, e26742.

Wolf, YI et al. (2016). “Two fundamentally different classes of microbial genes”. In: Nature microbiology 2, p. 16208.

Yao, X et al. (2015). “Isolation and characterization of a native avirulent strain of *Streptococcus suis* serotype 2: a perspective for vaccine development”. In: Sci Rep. 5, p. 9835.

Yun, KW et al. (2015). “Diversity of pneumolysin and pneumococcal histidine triad protein D of *Streptococcus pneumoniae* isolated from invasive diseases in korean children”. In: PLoS One 10.8, e0134055.

Zhang, A et al. (2011). “Comparative genomic analysis of Streptococcus suis reveals significant genomic diversity among different serotypes”. In: BMC Genomics 12, p. 523.

Zhang, Z et al. (2006). “KaKs_Calculator: calculating Ka and Ks through model selection and model averaging”. In: Genomics Proteomics Bioinformatics 4.4, pp. 259–263.

Zheng, WX et al. (2015). “Essentiality drives the orientation bias of bacterial genes in a continuous manner”. In: Scientific reports 5, p. 16431.

Zhou, CE et al. (2006). “MvirDB—a microbial database of protein toxins, virulence factors and antibiotic resistance genes for bio-defence applications”. In: Nucleic acids research 35.suppl_1, pp. D391–D394.

Zhou, Y et al. (2011). “PHAST: a fast phage search tool”. In: Nucleic acids research 39.suppl_2, W347–W352.

